## Supplementary Materials for "The Importance of Individual Beliefs in Assessing Treatment Efficacy: Insights from Neurostimulation Studies"

### Supplementary Information

Below, we report the model comparison for Study 1-5. In the column *Models*, we specify the term of the model that is added to test model fit. For example, in the second row, we test whether adding the interaction of *subjective treatment* and *time* to the baseline model (including the main effect of *time* and the interaction of *time* and *objective treatment*) leads to a better fit.

#### Study 1

First, we run model comparison considering depression scores over time as the outcome. Following, we compare models for remission and response rates, as well as anxiety scores.

**Table S1.**

*Count of Participants by Experimental Condition and Their Subjective Guess*

| Objective treatment | Subjective treatment | n |
|---|---|---|
| Bilateral rTMS | Subjective active | 25 |
| Bilateral rTMS | Subjective sham | 15 |
| Unilateral rTMS | Subjective active | 19 |
| Unilateral rTMS | Subjective sham | 21 |
| Sham rTMS | Subjective active | 11 |
| Sham rTMS | Subjective sham | 30 |

***Depression***

**Table S2.**

*Model comparison for the contribution of subjective treatment to HAMD-17 depression scores.*

| Models | Model | df | AIC | BIC | Test | L-ratio | p-value |
|---|---|---|---|---|---|---|---|
| *Time + Time: objective treatment* | 1 | 9 | 2028.28 | 2062.52 | | | |
| *Time: subjective treatment* | 2 | 11 | 1985.62 | 2027.48 | 1 vs 2 | 46.65 | **0.0000000001**$^{***}$ |
| *Time: subjective treatment: objective treatment* | 3 | 15 | 1988.50 | 2045.58 | 2 vs 3 | 5.11 | 0.2755 |

**Table S3.**

*Model comparison for the contribution of objective treatment to HAMD-17 depression scores.*

| Models | Model | df | AIC | BIC | Test | L-ratio | p-value |
|---|---|---|---|---|---|---|---|
| *Time+ Time: subjective treatment* | 1 | 7 | 1977.85 | 2004.49 | | | |
| *Time: objective treatment* | 2 | 11 | 1985.62 | 2027.47 | 1 vs 2 | 0.23 | 0.993 |
| *Time: subjective treatment: objective treatment* | 3 | 15 | 1988.50 | 2045.58 | 2 vs 3 | 5.11 | 0.275 |

**Table S4.**

*Model comparison for the contribution of subjective treatment to binary HAMD-17 response rates.*

| Models | df | Resid. Dev | Test | deviance | p-value |
|---|---|---|---|---|---|
| *Objective treatment* | 97 | 82.94 | | | |
| *Subjective treatment* | 96 | 68.15 | 1 vs 2 | 14.78 | **0.00012**$^{***}$ |
| *Subjective treatment: objective treatment* | 95 | 66.60 | 2 vs 3 | 1.55 | 0.410 |

**Table S5.**

*Model comparison for the contribution of objective treatment to HAMD-17 binary response rates*

| Models | df | Resid. Dev | Test | deviance | p-value |
|---|---|---|---|---|---|
| *Subjective treatment* | 97 | 69.27 | | | |
| *Objective treatment* | 95 | 67.86 | 1 vs 2 | .40 | 0.496 |
| *Subjective treatment: objective treatment* | 93 | 66.08 | 2 vs 3 | 1.78 | 0.410 |

**Table S6.**

*Model comparison for the contribution of subjective treatment to BDI-II binary response rates.*

| Models | df | Resid. Dev | Test | deviance | p-value |
|---|---|---|---|---|---|
| *Objective treatment* | 96 | 97.71 | | | |
| *Subjective treatment* | 95 | *86.90* | 1 vs 2 | 10.81 | **0.001**$^{***}$ |

| | | | | | |
|---|---|---|---|---|---|
| *Subjective treatment: objective treatment* | 93 | 86.27 | 2 vs 3 | 0.63 | 0.731 |

**Table S7.**

*Model comparison for the contribution of objective treatment to BDI-II binary response rates.*

| **Models** | **df** | **Resid. Dev** | **Test** | **deviance** | **p-value** |
|---|---|---|---|---|---|
| *Subjective treatment* | 96 | 87.17 | | | |
| *Objective treatment* | 95 | 86.90 | 1 vs 2 | 0.27 | 0.873 |
| *Subjective treatment: objective treatment* | 93 | 86.27 | 2 vs 3 | 0.62 | 0.731 |

***Anxiety***

In this study, anxiety was measured weekly as a secondary outcome alongside depression. Our results showed that *subjective treatment* did not contribute to a significantly better fit, neither as a main effect (*df*=18, *AIC*=1496.81, *BIC*=-698.75, *P*=.07) nor as part of an interaction with *objective treatment* (*df*=18, *AIC*=1520.46, *BIC*=-696.79, *P*=.561). Likewise, *objective treatment* did not contribute to a better model fit when compared to *subjective treatment* (*df*=18, *AIC*=1496.81, *BIC*=-698.75, *P*=.287)**.**

### Study 2

**Table S8.**

***Count of Participants by Experimental Condition and Their Subjective Guess***

| **Objective treatment** | **Subjective treatment** | **n** |
|---|---|---|
| Active rTMS | Subjective active | 18 |
| Active rTMS | Subjective sham | 7 |
| Sham rTMS | Subjective active | 15 |
| Sham rTMS | Subjective sham | 12 |

**Table S9.**

*Model comparison for the contribution of subjective treatment to HDRS-24 depression scores.*

| **Models** | **Model** | **df** | **AIC** | **BIC** | **Test** | **L-ratio** | **p-value** |
|---|---|---|---|---|---|---|---|
| *Time+ Time: objective treatment* | 1 | 11 | 1604.16 | 1642.85 | | | |
| *Time: subjective treatment* | 2 | 15 | 1606.97 | 1659.73 | 1 vs 2 | 5.18 | 0.268 |
| *Time: subjective treatment: objective treatment* | 3 | 19 | 1601.91 | 1563.91 | 2 vs 3 | 13.05 | **0.010*** |

**Table S10.**

*Model comparison for the contribution of objective treatment to HDRS-24 depression scores.*

| Models | Model | df | AIC | BIC | Test | L-ratio | p-value |
|---|---|---|---|---|---|---|---|
| *Time+ Time: subjective treatment* | 1 | 11 | 1605.64 | 1644.33 | | | |
| *Time: objective treatment* | 2 | 15 | 1606.97 | 1659.73 | 1 vs 2 | 6.67 | 0.154 |
| *Time: subjective treatment: objective treatment* | 3 | 19 | 1601.91 | 1668.74 | 2 vs 3 | 13.05 | **0.010*** |

**Table S11.**

*Model comparison for the contribution of subjective treatment to HDRS-24 binary remission rates.*

| Models | df | Resid. Dev | Test | deviance | p-value |
|---|---|---|---|---|---|
| *Objective treatment* | 48 | 49.05 | | | |
| *Subjective treatment* | 47 | 47.98 | 1 vs 2 | 1.07 | 0.301 |
| *Subjective treatment* objective treatment* | 46 | 43.51 | 2 vs 3 | 4.47 | **0.035*** |

**Table S12.**

*Model comparison for the contribution of objective treatment to HDRS-24 binary remission rates.*

| Models | df | Resid. Dev | df | deviance | p-value |
|---|---|---|---|---|---|
| *Subjective treatment* | 48 | 49.29 | | | |
| *Objective treatment* | 47 | 47.98 | 1 vs 2 | 1.31 | 0.253 |
| *Subjective treatment* objective treatment* | 46 | 43.51 | 2 vs 3 | 4.47 | **0.035**[*] |

**Table S13.**

*Model comparison for the contribution of subjective treatment to HDRS-24 binary response rates.*

| Models | df | Resid. Dev | Test | deviance | p-value |
|---|---|---|---|---|---|
| *Objective treatment* | 48 | 58.32 | | | |
| *Subjective treatment* | 47 | 57.97 | 1 vs 2 | 0.355 | 0.552 |
| *Subjective treatment* objective treatment* | 46 | 49.80 | 2 vs 3 | 8.172 | **0.004**[**] |

**Table S14.**

*Model comparison for the contribution of objective treatment to HDRS-24 binary response rates.*

| **Models** | **df** | **Resid. Dev** | **Test** | **deviance** | **p-value** |
|---|---|---|---|---|---|
| *Subjective treatment* | 48 | 59.10 | | | |
| *Objective treatment* | 47 | 57.97 | 1 vs 2 | 1.13 | 0.287 |
| *Subjective treatment* objective treatment* | 46 | 49.80 | 2 vs 3 | 8.17 | **0.004**** |

### Study 3

**Table S15.**

***Count of Participants by Experimental Condition and Their Subjective Guess***

| **Objective treatment** | **Subjective treatment** | **n** |
|---|---|---|
| Active tDCS | Subjective active | 16 |
| Active tDCS | Subjective sham | 9 |
| Sham tDCS | Subjective active | 13 |
| Sham tDCS | Subjective sham | 15 |

**Table S16.**

*Model comparison for the contribution of subjective treatment to CASRS-I inattention scores.*

| **Models**[1] | **Model** | **df** | **AIC** | **BIC** | **Test** | **L-ratio** | **p-value** |
|---|---|---|---|---|---|---|---|
| *Time+ Time: objective treatment* | 1 | 5 | 602.68 | 616.00 | | | |
| *Time: subjective treatment* | 2 | 6 | 593.80 | 609.78 | 1 vs 2 | 10.87 | **0.0009*** |
| *Time: subjective treatment: objective treatment* | 3 | 7 | 595.76 | 614.40 | 2 vs 3 | 0.046 | 0.829 |

**Table S17.**

*Model comparison for the contribution of objective treatment to CASRS-I inattention scores.*

| **Models**[1] | **Model** | **df** | **AIC** | **BIC** | **Test** | **L-ratio** | **p-value** |
|---|---|---|---|---|---|---|---|
| *Time+ Time: objective treatment* | 1 | 5 | 608.41 | 621.72 | | | |
| *Time: subjective treatment* | 2 | 6 | 593.80 | 609.78 | 1 vs 2 | 16.60 | **0.00004*** |
| *Time: subjective treatment: objective treatment* | 3 | 7 | 595.76 | 614.40 | 2 vs 3 | 0.046 | 0.829 |

### Study 4

**Table S18.**

***Count of Participants by Experimental Condition and Their Subjective Guess***

| **Objective treatment** | **Subjective treatment** | **n** |
|---|---|---|
| Anode 1mA | Subjective active | 22 |
| Anode 1mA | Subjective sham | 8 |
| Cathode 1mA | Subjective active | 16 |
| Cathode 1mA | Subjective sham | 14 |
| Cathode 1.5mA | Subjective active | 22 |
| Cathode 1.5mA | Subjective sham | 8 |
| Cathode 2mA | Subjective active | 17 |
| Cathode 2mA | Subjective sham | 13 |
| Sham | Subjective active | 9 |
| Sham | Subjective sham | 21 |

**Table S19.**

*Model comparison for the contribution of subjective treatment to mind-wandering scores.*

| Models | Model | df | AIC | BIC | Test | L-ratio | p-value |
|---|---|---|---|---|---|---|---|
| *Objective treatment* | 1 | 145 | 286.88 | 304.94 | | | |
| *Subjective treatment* | 2 | 144 | 284.72 | 305.81 | 1 vs 2 | 4.08 | **0.045*** |
| *Subjective treatment* objective treatment* | 3 | 140 | 286.61 | 319.74 | 2 vs 3 | 1.45 | 0.219 |

**Table S20.**

*Model comparison for the contribution of objective treatment to mind-wandering scores.*

| Models | Model | df | AIC | BIC | Test | L-ratio | p-value |
|---|---|---|---|---|---|---|---|
| *Subjective treatment* | 1 | 148 | 284.84 | 293.88 | | | |
| *Objective treatment* | 2 | 144 | 284.72 | 305.81 | 1 vs 2 | 8.11 | 0.093 |
| *Subjective treatment* objective treatment* | 2 | 140 | 286.61 | 319.74 | 2 vs 3 | 1.45 | 0.219 |

**Table S21.**

*Model comparison for the contribution of subjective dosage to mind-wandering scores.*

| Models | Model | df | AIC | BIC | Test | L-ratio | p-value |
|---|---|---|---|---|---|---|---|
| *Objective treatment* | 1 | 145 | 286.88 | 304.94 | | | |
| *Subjective dosage* | 2 | 142 | 282.90 | 310.01 | 1 vs 2 | 3.21 | **0.025***|
| *Subjective dosage* objective treatment* | 3 | 130 | 295.46 | 358.70 | 2 vs 3 | 0.85 | 0.590 |

**Table S22.**

*Model comparison for the contribution of objective treatment to mind-wandering scores.*

| Models | Model | df | AIC | BIC | Test | L-ratio | p-value |
|---|---|---|---|---|---|---|---|
| *Subjective dosage* | 1 | 146 | 282.94 | 297.88 | | | |
| *Objective treatment* | 2 | 142 | 282.01 | 310.81 | 1 vs 2 | 1.94 | 0.106 |
| *Subjective dosage* objective treatment* | 3 | 139 | 285.46 | 321.74 | 2 vs 3 | 1.14 | 0.332 |

### Study 5

We here apply our approach to experimental study that examined the effect of non-invasive brain stimulation (NIBS) on participants' working memory. In this case, we show that the inclusion of *subjective treatment* did not explained variability in experimental outcomes.

This study (Murphy et al., 2020) compared the effects of different NIBS techniques, namely anodal tDCS, tRNS + DC-offset, or sham stimulation over the left dorsolateral prefrontal cortex (DLPFC) on working memory (WM) performance and task-related EEG oscillatory activity in forty-nine healthy adults. Participants were allocated to receive either anodal tDCS ($N$=16), high-frequency tRNS + DC-offset ($N$=16), or sham stimulation ($N$=17) to the left DLPFC using a between-subjects design. The Sternberg WM task was used to measure changes in WM performance before, 5-min and 25-min post-stimulation. Moreover, event-related synchronisation/desynchronization (ERS/ERD) of oscillatory activity was analysed from EEG recorded during WM encoding and maintenance. At the end of the experiment, participants were asked whether they thought they received active or sham stimulation (presented as a binary choice) via a short questionnaire.

We employed the same analytical approach as for Experiments 1-4, looking at the contribution of *subjective treatment* to model fit over and beyond *objective treatment* and examining whether *objective treatment* explained variability in experimental outcomes (WM accuracy and reaction times) over and beyond *subjective treatment*.

A linear mixed model with WM reaction time was fitted to the data for time 0-2 (before NIBS, 5 minutes after, and 25 minutes after). The baseline model included *objective treatment* (sham/tRNS/tDCS) as the main effect, as well as the interaction of *time* and *objective treatment*. We first added to this model *subjective treatment* (active or placebo) as a main effect. Next, we extended the model to include the two-way interaction of *time* and *subjective treatment*. Lastly, we considered a model with the three-way interaction of *time*, *subjective treatment* and *objective treatment*. Our results showed that the inclusion of *subjective treatment* did not lead to a better model fit neither as a main effect (*df*=13, *BIC*=1824.16, *AIC*=1785.28, *P*=0.670) nor as a two-way interaction with *time* (*df*=15, *BIC*=1833.25, *AIC*=1788.40, *P*=0.642) and a three-way interaction with *time* and *objective treatment* (*df*=21, *BIC*=1856.48, *AIC*=1793.70, *P*=0.348). Hence, participants' subjective experience about the treatment did not explain variability in reaction times beyond the actual treatment condition to which participants were assigned.

We next examined whether the changes in reaction time were explained by both *objective treatment*. We, therefore, added *objective treatment* first and, following, *objective treatment*time* after *subjective treatment*time* was already included in the baseline model. Our results showed that the inclusion of neither *objective treatment* (*df*=11, *BIC*=1820.58, *AIC*=1787.69, *P*=0.139) nor *objective treatment*time* (*df*=15, *BIC*=1833.25, *AIC*=1788.40, *P*=0.121) led to a better model fit. Therefore, when accounting for participants' subjective experience of receiving the real or placebo treatment, the actual treatment to which participants were assigned during the experiment did not contribute to explaining changes in WM reaction time.

Following, we run the same analysis with WM accuracy as the outcome of interest. Model comparison showed that *subjective treatment* did not lead to a better model fit neither as a main effect (*df*=13, *BIC*=1084.64, *AIC*=1045.77, *P*=0.069) nor as a two-way interaction with *time* (*df*=15, *BIC*=1094.60, *AIC*=1049.74, *P*=0.990) and a three-way interaction with *time* and *objective treatment* (*df*=21, *BIC*=1122.48, *AIC*=1059.68, *P*=0.914). On the contrary, we next examined whether the changes in WM accuracy were explained by *objective treatment*. Our results showed that the inclusion of *objective treatment* (*df*=11, *BIC*=1089.73, *AIC*=1056.84, *P*=0.183) did not lead to a better model fit, but the interaction of *objective treatment*time* did (*df*=15, *BIC*=1094.60, *AIC*=1049.74, *P*=0.005). Therefore, when accounting for participants' subjective experience of receiving the real or placebo treatment, the actual treatment to which participants were assigned during the experiment contributed to explaining changes in WM accuracy.
